# Functional interplay between SWI/SNF complexes underlies BRD9 dependency in SMARCB1-mutant cancers

**DOI:** 10.1101/2022.08.07.503080

**Authors:** Robert J. Mobley, Jacquelyn A. Myers, Kendall M. Wills, Hayden A. Malone, Trishabelle I. Manzano, Janet F. Partridge, Charles W. M. Roberts

## Abstract

Genes encoding subunits of SWI/SNF (BAF) chromatin remodeling complexes are mutated in >20% of cancers. SWI/SNF complexes exist in three distinct families that each contribute to regulation of transcription, although the functional interactions between the families are not well understood. Rhabdoid tumors constitute an informative model system as these highly aggressive cancers are driven by inactivation of a single SWI/SNF subunit, SMARCB1, which is present in two SWI/SNF families (cBAF and PBAF) but not in the third (GBAF/ncBAF). We and others have shown that BRD9, a therapeutically targetable member of ncBAF, is essential specifically in SMARCB1-deficient cancers, suggesting key functional relationships between SMARCB1-containing complexes and BRD9/ncBAF. However, the mechanistic underpinnings of these relationships are poorly understood. Here, we demonstrate that genomic binding of BRD9 is largely dependent upon SMARCB1 such that the absence of SMARCB1 results in significantly reduced BRD9 binding. At select sites, however, we show that SMARCB1-loss results in gain of BRD9 binding and BRD9-dependent accessibility. We find that this gain is associated with expression of genes promoting cell migration. Our results define relationships between SWI/SNF complex families, elucidate mechanisms by which SMARCB1 loss drives oncogenesis, and provide mechanistic insight into the synthetic-lethal relationship between SMARCB1 and BRD9.

## Introduction

Genes encoding subunits of SWI/SNF (BAF) chromatin remodeling complexes are mutated in >20% of all human cancers (Kadoch et al., 2013; Orlando et al., 2019; Shain & Pollack, 2013). SWI/SNF complexes hydrolyze ATP to mobilize nucleosomes and in cooperation with transcription factors are critical for dynamic management of genomic accessibility and gene expression (Alver et al., 2017; Brien et al., 2018; Iurlaro et al., 2021; Langer et al., 2019; Mashtalir et al., 2021; Nakayama et al., 2017; X. Wang et al., 2017). The three families of SWI/SNF complexes contain numerous shared subunits but also subunits that are exclusive to specific families. Two of the families (cBAF and PBAF) contain the nucleosome-interacting subunit SMARCB1, but the third (GBAF/ncBAF) does not (Alpsoy & Dykhuizen, 2018; Mashtalir et al., 2018). All three families contribute to transcriptional regulation, but the relationship between the families, including whether they cooperate or antagonize each other, is poorly understood. Here, we sought to identify the different contributions of SMARCB1-containing complexes and BRD9-containing complexes to the regulation of chromatin organization and gene expression.

SMARCB1 was implicated as a tumor suppressor when it was shown that nearly all cases of rhabdoid tumors (RTs), a highly lethal cancer of early childhood, contain biallelic inactivation of SMARCB1 as the sole recurrent mutation (Biegel et al., 1999; Versteege et al., 1998). Subsequently, compromising mutations in SMARCB1 were identified in several types of pediatric and adult cancers, as well as in neurodevelopmental disorders (Holsten et al., 2018). Deletion of SMARCB1 in mice results in a fully penetrant cancer phenotype with a median onset of only 11 weeks thus demonstrating a bona fide role for SMARCB1 as a tumor suppressor (Guidi et al., 2001; Klochendler-Yeivin et al., 2000; Roberts et al., 2000, 2002). The mechanism of tumor suppressor activity was later illuminated when it was shown that SMARCB1 enables SWI/SNF activation of enhancers that are required for lineage specification such that loss of SMARCB1 impairs differentiation (Alver et al., 2017; Nakayama et al., 2017; X. Wang et al., 2017).

Recently, SMARCB1-deficient cancer cells were demonstrated to specifically depend upon BRD9 for survival, demonstrating a clear functional relationship between these mutually exclusive SWI/SNF complex members (Brien et al., 2018; Michel et al., 2018; X. Wang et al., 2019). Additionally, the SMARCB1-containing cBAF and PBAF complexes (hereafter referred to as SMARCB1-BAF complexes) and the BRD9-containing ncBAF complex have recently been shown to have distinct affinities for, and to be variably activated by, specific covalent modifications of histone residues (Mashtalir et al., 2021). These findings elevate important unaddressed questions about chromatin management in development and disease: What are the distinct contributions of SMARCB1-BAF and ncBAF complexes in regulation of genomic accessibility and gene expression? And does SMARCB1-loss result in altered gene regulatory roles of BRD9-containing ncBAF complexes?

Here, we investigate the interplay of BRD9- and SMARCB1-BAF in gene regulation by using SMARCB1-deficient RT cells engineered to rapidly induce expression of SMARCB1 as well as taking advantage of dBRD9-A, a degrader compound that rapidly and near-completely degrades BRD9 (Brien et al., 2018; Garvin et al., 1993). Together, our data reveal how SMARCB1-loss results in aberrant control of proliferation and migration by ncBAF complexes. This work advances our understanding of the relationships between SMARCB1 and BRD9-containing SWI/SNF complexes and the roles BRD9 assumes in the absence of SMARCB1.

## Results

### BRD9 regulates genomic accessibility and gene expression in the absence of SMARCB1

To determine the interplay between SMARCB1-containing and BRD9-containing SWI/SNF complexes in gene regulation, we engineered SMARCB1-deficient G401 RT cells to induce SMARCB1 expression and used a BRD9 degrader (dBRD9-A) to rapidly deplete BRD9 protein from both the SMARCB1-induced and control cells (Brien et al., 2018). After 24 h of SMARCB1 induction, we saw robust and reproducible levels of SMARCB1 protein (Figure 1A). Glycerol gradient fractionation revealed that SMARCB1 was effectively integrated into SMARCA4-containing SMARCB1-BAF complexes without affecting existing levels of ncBAF complexes (Figure 1B). Further, 24 h of treatment with dBRD9-A resulted in near-total depletion of BRD9 protein (Brien et al., 2018) (Figure 1A). Having thus validated our experimental system, we began to examine the relative contributions of SMARCB1- and BRD9-containing SWI/SNF complexes in the creation of genomic accessibility and control of gene expression.

**Figure 1:**
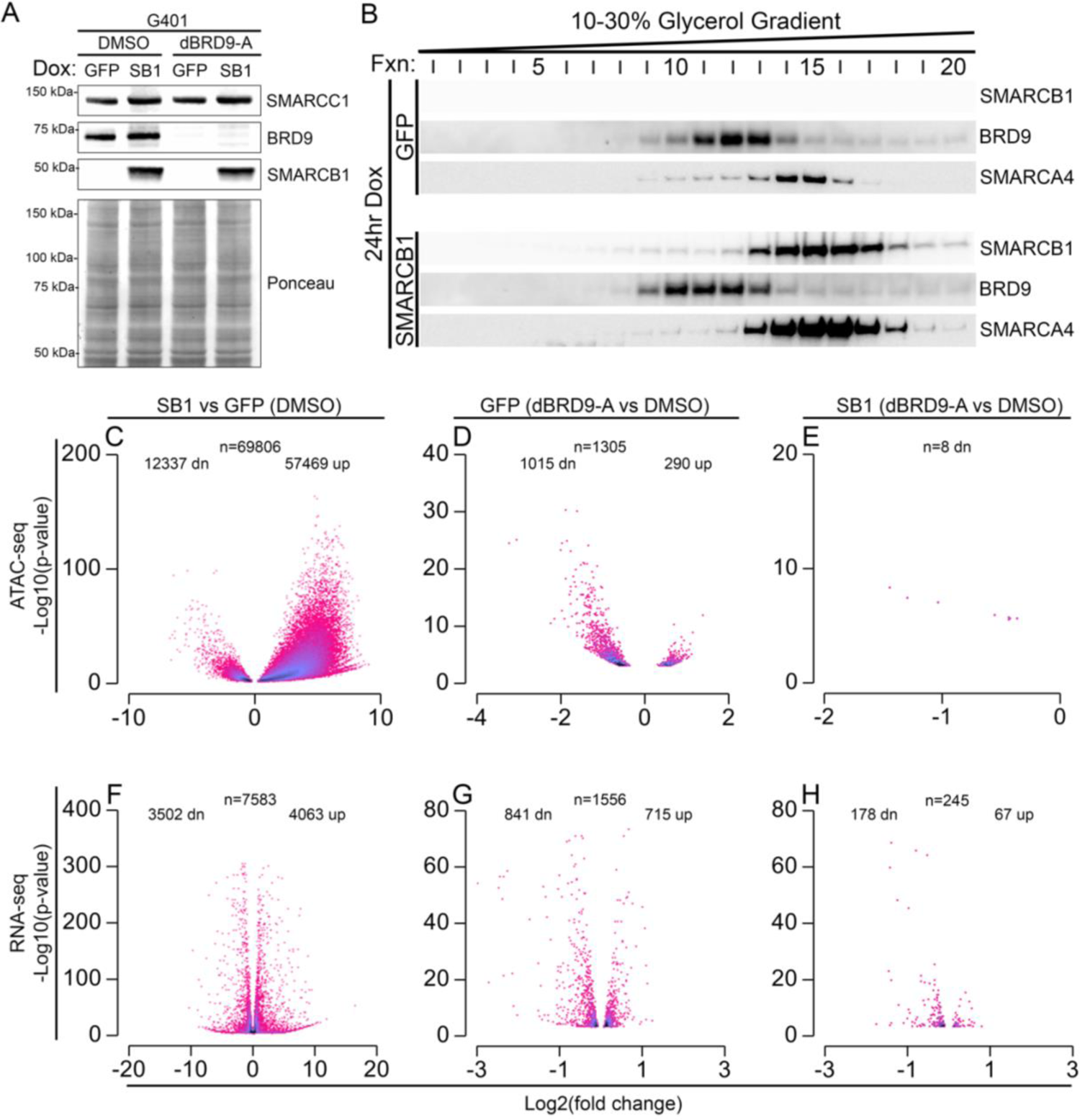
BRD9 gains gene-regulatory roles in the absence of SMARCB1, a key regulator of genomic accessibility and gene expression. A. Immunoblot demonstrating SMARCB1 expression following 24-h doxycycline induction of SMARCB1 (SB1 vs. GFP) and BRD9 degradation following 24-h treatment with 100 nM dBRD9-A (dBRD9-A vs. DMSO) in G401 RT cells. SMARCC1 levels were unaffected by either treatment. Images shown are representative of 3 independent experiments. Ponceau stain is shown as the protein loading control. B. Immunoblot of glycerol gradient fractionation showing density sedimentation of SMARCB1, BRD9, and ATPase subunit SMARCA4 after 24-h SMARCB1-rescue (SB1) or control (GFP) induction. Images shown are representative of 2 independent experiments. C-E. Volcano plots displaying ATAC-seq data comparing SMARCB1 (SB1)-induced vs. GFP-induced G401 cells (C), dBRD9-A-treated vs. DMSO-treated GFP-induced cells (D), and dBRD9-A-treated vs. DMSO-treated SB1-induced cells (E), treated as described in (A). Data shown are the average of 3 biological replicates, FDR < 0.05. F-H. Volcano plots displaying RNA-seq data comparing SMARCB1 (SB1)-induced vs. GFP-induced G401 cells (F), dBRD9-A-treated vs. DMSO-treated GFP-induced cells (G), and dBRD9-A-induced vs. DMSO-treated SB1-induced cells (H), treated as described in (A). Data shown are the average of 3 biological replicates, p <0.001.

We used ATAC-seq to evaluate the effects of SMARCB1 and BRD9 directed nucleosome remodeling activity. SMARCB1 has previously been shown to facilitate enhancer activation and associated gene expression (Alver et al., 2017; Nakayama et al., 2017; X. Wang et al., 2017). Accordingly, SMARCB1 rescue resulted in robust gain of genomic accessibility (57,469 sites, Figure 1C, FDR < 0.05). We also detected a substantial number of sites with decreased genomic accessibility (12,337 sites, Figure 1C, FDR < 0.05), raising the possibility that SMARCB1 could also play a role in limiting genomic accessibility. In contrast, degradation of BRD9 resulted in far fewer changes than expression of SMARCB1. In SMARCB1-deficient G401 cells, degradation of BRD9 resulted in 1,015 significant decreases and 290 increases in accessibility (Figure 1D, FDR < 0.05). In SMARCB1-induced cells, degradation of BRD9 caused only 8 significant decreases in accessibility (Figure 1E, FDR < 0.05). Together, these data suggest that SMARCB1 regulates genomic accessibility at more sites than BRD9, and that the loss of SMARCB1 results in select sites becoming dependent upon the BRD9-containing ncBAF complex for genomic accessibility.

Next, we asked whether these chromatin accessibility changes resulted in altered gene expression. Indeed, SMARCB1-rescue led to upregulation of 4,063 and downregulation of 3,502 genes, whereas degradation of BRD9 in SMARCB1-deficient G401 cells resulted in 715 upregulated and 843 downregulated genes (Figure 1 F-G, p < 0.001). As seen in the ATAC studies, BRD9 degradation in SMARCB1-induced cells showed minimal effects, with only 67 upregulated genes and 178 downregulated genes (Figure 1H, p < 0.001). Together these data suggest that SMARCB1-containing SWI/SNF complexes have much greater roles in the control of genomic accessibility and gene expression than do the mutually exclusive BRD9-containing complexes. However, in the absence of SMARCB1, as occurs in RT, BRD9-containing complexes gain regulatory roles controlling accessibility and gene expression at some sites.

### SMARCB1-rescue redistributes BRD9 genomic binding

We next sought to further understand how the loss of SMARCB1 affects ncBAF complexes by evaluating the consequences of SMARCB1 re-expression in RT cells. Given that cells induced to express SMARCB1 showed reduced ncBAF-mediated control of genomic accessibility, we asked whether SMARCB1-BAF directly displaces BRD9 from these sites. Using CUT and RUN, we generated genomic binding profiles of BRD9 in G401 RT cells both in the absence and presence of SMARCB1 and compared these profiles with SMARCB1 genomic binding data. To ensure that we were monitoring binding of SWI/SNF complexes, we integrated BRD9 and SMARCB1 binding data with that obtained for SMARCA4, the sole SWI/SNF ATPase subunit expressed in G401 cells. Additionally, to associate sites bound by SMARCB1 or BRD9 with promoters and enhancers, we performed MNase-ChIP for H3K27Ac, H3K4me1, and H3K4me3 histone modifications in SMARCB1-vs. control (GFP)-induced cells.

First, we analyzed SMARCB1-bound sites. CUT and RUN genomic profiling methods enabled high-resolution detection of SMARCB1 binding, yielding results consistent with previous ChIP studies (Nakayama et al., 2017; X. Wang et al., 2017). After 24 h of SMARCB1 induction, we detected 94,114 SMARCB1-bound loci (Figure 2A), which were predominantly associated with enhancers (H3K27Ac+), primed enhancers (H3K27Ac-, H3K4me1+), and promoters (+/-1kb of TSS, H3K4me3+). Using H3K27Ac ChIP-seq data from SMARCB1-induced G401 cells and the Rank Ordering of Super Enhancers (ROSE) algorithm, we determined that 19% of SMARCB1-bound sites were within traditional enhancers (TEs), and 17% were within super enhancers (SEs) (Lovén et al., 2013; Whyte et al., 2013), both of which also gained enrichment of SMARCA4 and the enhancer-associated histone modification H3K4me1 upon SMARCB1 expression (Figure 2A). Only 6% of SMARCB1 peaks were found at promoters (Figure 2A). Most SMARCB1-binding sites (55,322; 59%) were detected more than 1 kb from a TSS and were marked by H3K4me1 but not H3K27Ac (Figure 2A). We designated these sites as primed enhancers (PEs) (Figure 2A). These results show that SMARCB1 is predominantly recruited to regions of the genome that are either primed or active enhancers.

**Figure 2:**
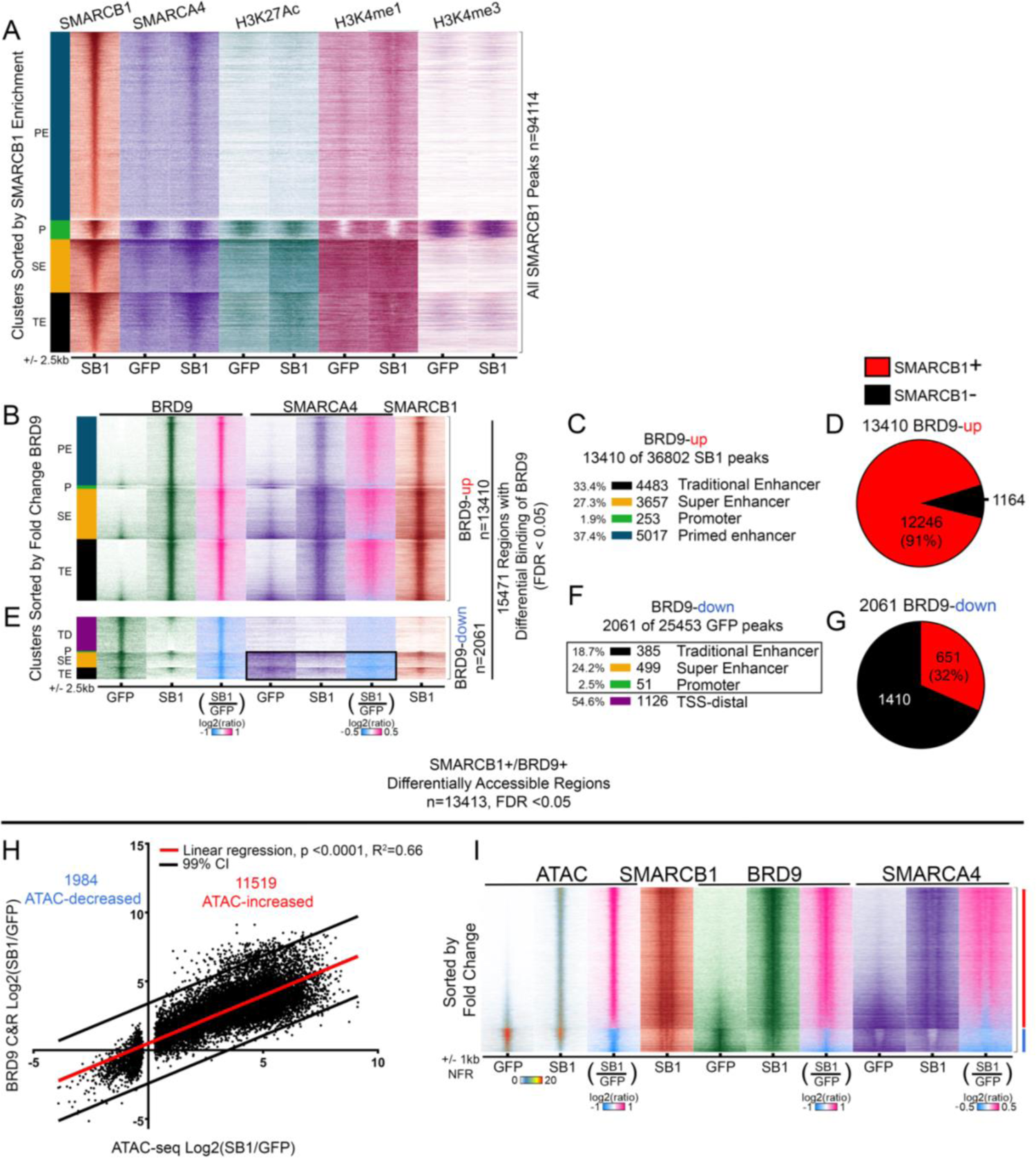
SMARCB1-rescue redistributes BRD9 genomic binding. A. SMARCB1 CUT & RUN following expression of SMARCB1 in G401 RT cells for 24 h identifies 94,114 SMARCB1 peaks. Genomic binding signal from SMARCB1- and GFP-induced G401 (SB1, GFP) cells is sorted by SMARCB1 enrichment and clustered by genomic elements defined by chromatin marks generated from MNase-ChIP. Traditional enhancers (TEs, black) and super enhancers (SEs, gold) were called by using H3K27Ac data from SB1 cells according to Rank Ordering of Super Enhancers (ROSE). Promoter-associated peaks outside a ROSE-defined enhancer (Ps, green) were called within 1 kb of an H3K4me3-enriched promoter. Primed enhancer sites (PEs, dark blue) are H3K4me1-enriched, H3K27Ac-negative regions >1 kb from the nearest TSS. Data shown are the average of two biological replicates. B-G. BRD9, SMARCA4, and SMARCB1 genomic binding data from 15,471 differentially bound BRD9 CUT & RUN peaks (SB1 vs. GFP, FDR < 0.05). BRD9 CUT & RUN data are the average of 3 biological replicates. B-D BRD9, SMARCA4, and SMARCB1 genomic binding data from 13,410 sites with significant increase in BRD9 upon SMARCB1 expression (SB1 vs. GFP, FDR < 0.05) (B). Sites gaining BRD9 are annotated with ROSE and histone modification data analysis from SMARCB1-induced G401 cells (C). Circle graphs show the percentage of BRD9-up sites bound by SMARCB1 (D). E-G. BRD9, SMARCA4, and SMARCB1 genomic binding data from 2,061 sites with a significant decrease in BRD9 (SB1 vs. GFP, FDR < 0.05) (E). Sites losing BRD9 are annotated with ROSE and histone modification data analysis from GFP-induced (negative control) G401 cells (F). Sites >1 kb from a TSS with no H3K4me1 were classified as TSS-distal sites (TDs). A black box indicates a region within TEs, SEs, and Ps where SMARCA4 is also lost upon SMARCB1 rescue (E, F). Circle graphs show the percentage of BRD9-down sites bound by SMARCB1 (G). H. Linear regression comparing SMARCB1-dependent changes in accessibility to fold-change BRD9 enrichment between SMARCB1 (SB1)- and GFP-induced G401 cells at sites where SMARCB1 and BRD9 were detected in at least one sample at a differentially accessible region (SMARCB1^+^/BRD9^+^ Differentially Accessible Regions). SMARCB1^+^/BRD9^+^ differentially accessible regions are correlated with SMARCB1-dependent changes in BRD9 binding (p<0.0001, R^2^=0.66). I. Genomic heatmaps of ATAC-seq data, SMARCB1 and BRD9 CUT & RUN, and SMARCA4 ChIP-seq at the sites from (H) that are bound by SMARCB1 and BRD9 that show SMARCB1-dependent changes in ATAC accessibility, centered on the differentially accessible nucleosome-free regions. Sites are sorted by difference in the ATAC-seq nucleosome-free signal measured between SMARCB1 (SB1)- and GFP-induced G401 cells. The red sidebar indicates ATAC-up sites. The blue sidebar indicates ATAC-down sites.

We then asked how SMARCB1 rescue affected BRD9 binding. The number of BRD9 peaks increased following SMARCB1 rescue (29,504 BRD9 peaks vs. 21,510 BRD9 peaks) which were enriched within enhancers and promoters (Figure 2 B-D, Supplemental Figure 1 A-D). To specifically characterize SMARCB1-dependent changes in BRD9 genomic binding, we focused on sites where SMARCB1 rescue resulted in significantly increased (n=13,410, BRD9-up) or decreased (n=2061, BRD9-down) BRD9 binding (Figure 2 B, E). Upon SMARCB1 rescue, BRD9 was preferentially gained at enhancers (PEs, TEs, SEs). SMARCA4 was also gained at these sites, suggesting that BRD9 was not binding alone but was likely present within ncBAF complexes (Figure 2 B, C). SMARCB1 was bound at 91% of these sites, suggesting that SMARCB1-containing complexes facilitate the binding of ncBAF (Figure 2D). Sites that lost BRD9 upon SMARCB1 rescue were frequently at enhancers (TEs and SEs) but were also at non-enhancer–marked TSS-distal sites (TSS-distal) (Figure 2E, F). SMARCB1 was bound at nearly one-third of the sites that lost BRD9, suggesting that SMARCB1 antagonizes BRD9 binding at these sites (32%, Figure 2 E, G). Sites displaying SMARCB1-BRD9 antagonism were predominantly at traditional enhancers and super enhancers (Figure 2 E, F) and SMARCA4 was reduced at these sites on SMARCB1 rescue, indicating that overall levels of SWI/SNF binding were somewhat decreased. In contrast, SMARCA4 was never detected at the non-enhancer TSS-distal sites that lost BRD9, suggesting that a substantial portion of BRD9 binding events in SMARCB1-deficient cells were independent of the ncBAF complex (Figure 2 E).

We hypothesized that SMARCB1 might be required to generate accessibility for BRD9 binding because SMARCB1-rescue resulted in many increases in genomic accessibility and BRD9 binding. To test this notion, we examined changes in BRD9 binding at SMARCB1-bound sites that showed significant changes in genomic accessibility (Supplemental Figure 2E, Figure 2H, I). Regions of SMARCB1-promoted genomic accessibility were strongly correlated with enhanced BRD9 binding at these sites, and the converse was also true (Figure 2H, I). These data suggest that changes in BRD9 genomic binding are largely determined by SMARCB1-dependent management of genomic accessibility.

### BRD9 and SMARCB1 have distinct patterns of interaction with nucleosome-free regions

We next asked how SMARCB1-regulated genomic accessibility could direct both gains and losses of BRD9. We examined whether the pattern of SMARCB1 enrichment at sites of decreased accessibility differed from that at sites of increased accessibility. SMARCB1 directly anchors SWI/SNF to the nucleosome acidic patch (Valencia et al., 2019), so we hypothesized that SMARCB1 would be preferentially enriched on nucleosomes that flank newly accessible regions. Since BRD9 binding was largely determined by SMARCB1, we asked whether BRD9 would bind to nucleosome-free regions or to nucleosomes that flank accessible regions managed by SMARCB1.

To determine the localization of SMARCB1 and BRD9 with respect to nucleosome occupancy, we utilized SMARCB1 and BRD9 CUT & RUN and ATAC-seq data and restricted our analyses to SMARCB1 and BRD9-bound loci that changed accessibility upon SMARCB1-rescue. We detected sites that gained ATAC accessibility (Figure 3A-D) and those that lost ATAC accessibility (Figure 3E-H). Specifically at sites that gained ATAC accessibility, we detected *de novo* gain of signal from flanking nucleosomes (Figure 3 A) and nucleosome-free (Figure 3 B) regions. BRD9 binding was focused on the nucleosome-free region while SMARCB1 and SMARCA4 showed a bi-modal distribution at these same sites, with highest enrichment at the flanking nucleosomes (Figure 3 C, D), indicating that SMARCB1-BAF interacts with the flanking nucleosomes to generate accessible regions that then recruit BRD9. Thus SMARCB1-BAF maintains genomic accessibility and enables ncBAF binding at traditional, super, and primed enhancer regions (Supplemental Figure 2 A). In contrast, at sites where SMARCB1-rescue led to decreased ATAC accessibility (Figure 3 E-H), in addition to binding the flanking nucleosomes, SMARCB1 also occupied the nucleosome-free space. Although BRD9 and SMARCA4 binding were both decreased at the ATAC-decreased sites, their overall binding pattern was similar at ATAC-increased and -decreased sites (Figure 3 A-D vs E-H). Notably, sites that lost ATAC accessibility on SMARCB1-rescue showed exceptionally high accessibility in the absence of SMARCB1 and were bound by BRD9. Upon SMARCB1-rescue, this accessibility was decreased, but was still higher than at sites that gained *de novo* accessibility after SMARCB1-rescue (Figure 3 A, B vs. E, F). These data suggest that SMARCB1-loss results in specific regions gaining “hyper-accessibility” and BRD9 binding, thereby restructuring chromatin for the expression of aberrant transcriptional programs seen in SMARCB1-deficient cancers.

**Figure 3:**
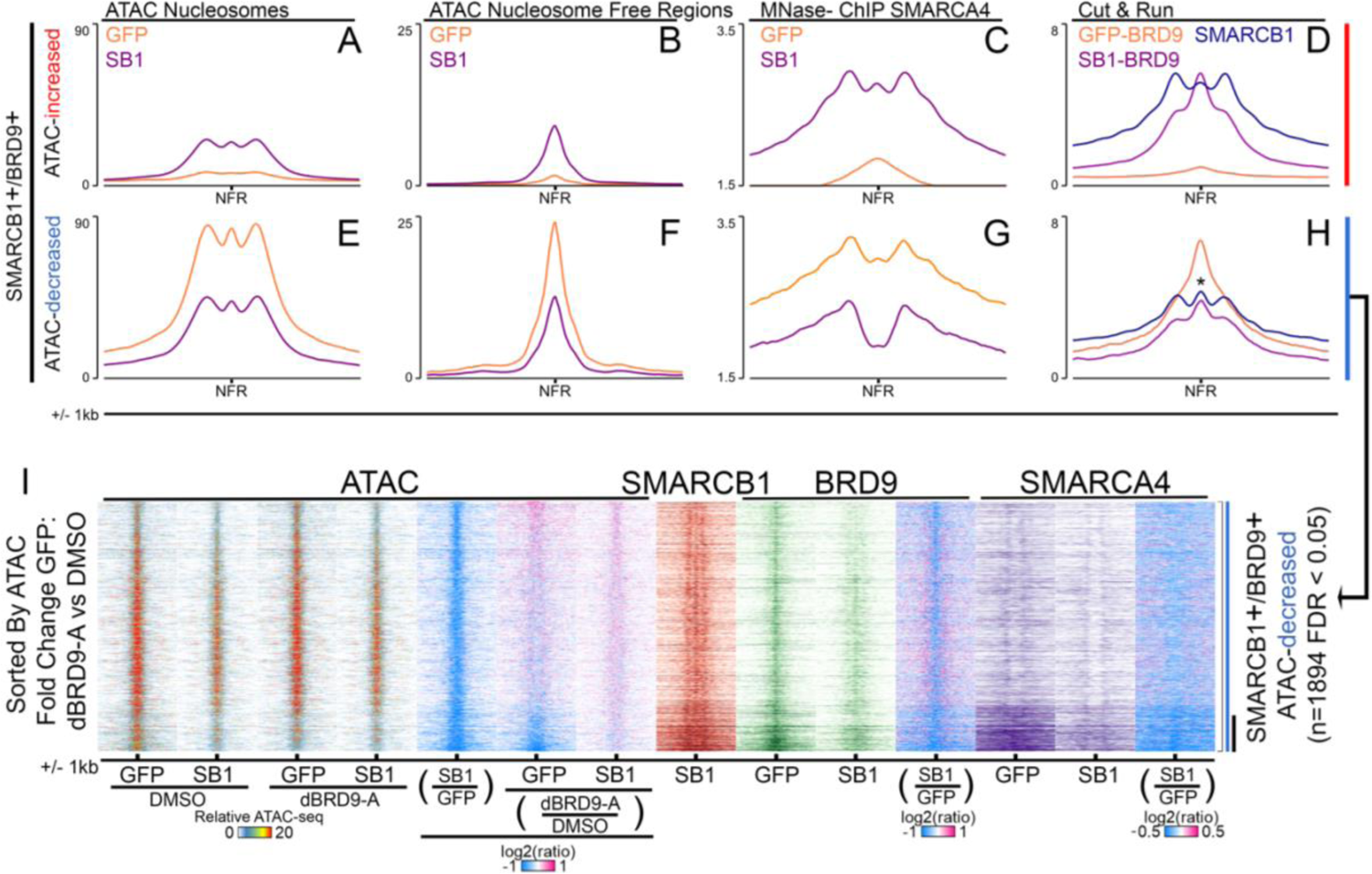
Functional interplay between SMARCB1 and BRD9 at sites with SMARCB1-dependent change in accessibility. A-D. Metaplots of nucleosome positioning (A) and nucleosome-free regions (B) derived from ATAC-seq data, SMARCA4 ChIP (C), and SMARCB1 and BRD9 Cut & Run data (D) at SMARCB1^+^/BRD9^+^ differentially accessible regions where ATAC signal is increased upon SMARCB1- or GFP induction in G401 cells (SB1, GFP) (sites with red sidebar in Figure 2I). E-H. Metaplots of nucleosome positioning (E) and nucleosome-free regions (F) derived from ATAC-seq data, SMARCA4 MNase-ChIP (G), and SMARCB1 and BRD9 Cut & Run data (H) at SMARCB1^+^/BRD9^+^ differentially accessible regions where ATAC signal is decreased upon SMARCB1- or GFP induction in G401 cells (SB1, GFP) (sites with blue sidebar in Figure 2I). I. Genomic heatmaps of SMARCB1^+^/BRD9^+^ differentially accessible regions where ATAC signal is decreased upon SMARCB1-rescue from E-H and Figure 2I, compared with ATAC-seq data from SMARCB1- or GFP-induced G401 cells (SB1, GFP) treated with dBRD9-A or DMSO for 24 h. At the subset of sites where BRD9 loss also decreases accessibility (black sidebar), SMARCB1 rescue partially depletes BRD9 and SMARCA4.

We then asked whether sites whose genomic accessibility was regulated by SMARCB1 were associated with specific gene regulatory elements and therefore likely to hold disease relevance in SMARCB1-deficient cancers. ATAC-increases were almost exclusively at enhancers (traditional, primed, and super), whereas ATAC-decreases were more often found at promoters and non-enhancer TSS-distal sites (Supplemental Figure 3 A-B). Overall, these data suggest that the roles of SMARCB1 include distinct modes of generation and limitation of genomic accessibility that can promote or prevent BRD9 binding at specific gene regulatory elements.

### ncBAF and SMARCB1-BAF have opposing roles at select enhancer sites

SMARCB1-deficient cells depend upon BRD9 for viability. We therefore asked whether this dependence could be due to BRD9 promoting hyper-accessibility in SMARCB1-deficient cells at the sites described above that lose both binding of BRD9 and ATAC accessibility upon SMARCB1-rescue. To address this question, we degraded BRD9 in RT cells that lack SMARCB1 and performed ATAC-seq. Following BRD9 degradation, most of the sites that lost accessibility upon SMARCB1-rescue were unaffected (Figure 3I, ATAC-GFP Log2(dBRD9-A/DMSO)), but a subset (237 of the 1894 sites) lost accessibility (Figure 3I, marked by black bar). Whilst sites that were unaffected by loss of BRD9 included promoters and TSS-distal elements, sites that lost accessibility upon either BRD9 degradation or SMARCB1 rescue were rarely near promoters but were almost exclusively within traditional and super enhancers (Supplemental Figure 2B). As expected (Figure 1E), degradation of BRD9 in RT cells expressing SMARCB1 did not affect accessibility of these enhancer regions (Figure 3I). Thus, at sites where SMARCB1-rescue dampens chromatin accessibility, BRD9 is largely dispensable in the presence of SMARCB1. However, in cells lacking SMARCB1, BRD9 maintains hyper-accessibility at a subset of traditional and super-enhancer loci (Figure 3I). Together, these data suggest that SMARCB1-BAF and ncBAF can hold opposing roles in enhancer regulation at select sites, demonstrating a functional difference between SMARCB1- and BRD9-containing SWI/SNF complexes that may underly BRD9-dependency in SMARCB1-deficient cancer.

### BRD9 supports overexpression of genes driving G401 cell migration

Considering the above findings, we next sought to evaluate the consequences of SMARCB1 and BRD9 interplay in the control of transcriptional regulation. To this end, we used Binding and Expression Target Analysis (BETA) to associate changes in ATAC-seq with specific changes in gene expression (Supplemental Figure 3 A-D)(S. Wang et al., 2013). Sites where both SMARCB1 and BRD9 were directly bound and that showed enhanced accessibility upon expression of SMARCB1 were significantly associated with 1541 upregulated genes (Figure 3A-D, Supplemental Figure 4A, Supplemental File 1). In contrast, sites where SMARCB1 and BRD9 were both bound but that showed reduced accessibility upon SMARCB1 expression (Figure 3 E-H) were significantly associated with 1000 downregulated genes (Supplemental Figure 4B, Supplemental File 2). Collectively these data demonstrate that loss of SMARCB1, as occurs in RT, predominantly results in loss of BRD9 binding, reduced accessibility at enhancers, and downregulation of associated genes. However, at a subset of sites, SMARCB1 loss results in increased BRD9 binding that correlates with enhanced accessibility and transcriptional upregulation.

Given the correlation between BRD9 binding and enhanced accessibility at a subset of sites we next sought to evaluate causation. Specifically, we sought to determine whether it was the binding of BRD9 that drove transcriptional upregulation. Therefore, we asked whether BRD9-dependent genomic accessibility contributed to transcriptional regulation. In cells lacking SMARCB1, degradation of BRD9 resulted in loss of genomic accessibility associated with significant downregulation of 307 genes (p= 3.22 × 10^-11^) (Supplemental Figure 3C-D, Supplemental File 3). Given our hypothesis that SMARCB1 rescue also depleted BRD9 from specific sites, we asked if there was overlap between genes downregulated following SMARCB1-rescue and those downregulated following BRD9 degradation. Of the 307 BRD9-dependent genes, 98 were also associated with regions that showed decreased genomic accessibility upon SMARCB1 rescue (Figure 3I). Together these data suggest that SMARCB1 limits expression of some BRD9-dependent genes (98) and that SMARCB1-loss permits their BRD9-dependent overexpression.

We next evaluated how genes that are activated in the absence of SMARCB1 might be supporting the cancer phenotype of G401 RT cells. We performed gene set enrichment analysis in GO: Biological Function categories using Enrichr (Chen et al., 2013; Kuleshov et al., 2016; Xie et al., 2021) and found that these genes were most significantly enriched in categories impacting translation (Figure 4A). Specifically, ribosomal subunit and ribosomal/non-coding RNA gene sets were downregulated after SMARCB1-rescue (Figure 4A). These data are of interest given our previously published work demonstrating that RT are specifically vulnerable to translational inhibitors (Howard et al., 2020).

**Figure 4:**
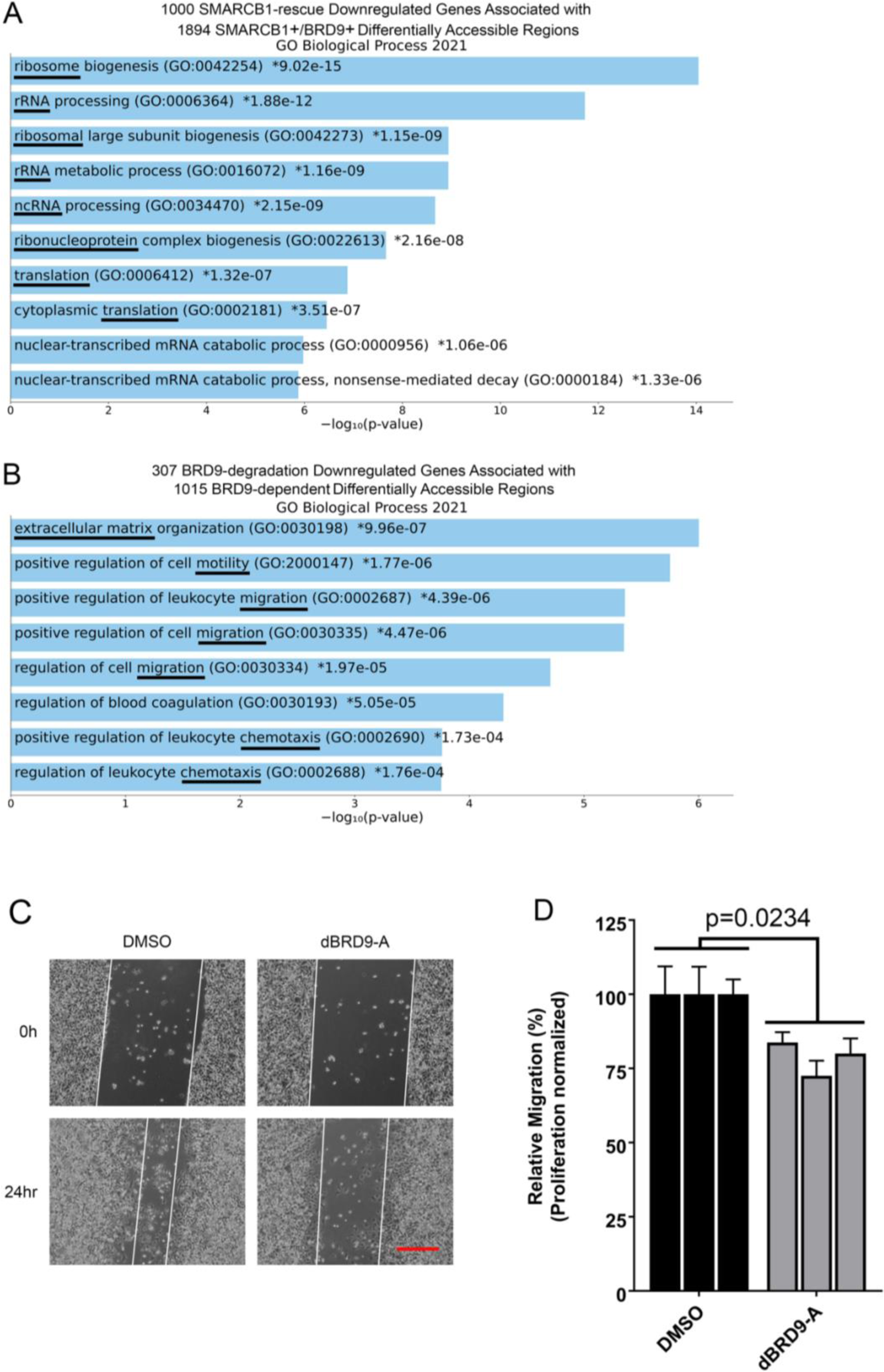
BRD9 supports overexpression of genes in SMARCB1-deficient G401 RT cells. A. GSEA of 1000 downregulated genes that are associated with SMARCB1-bound sites that show decreased genomic accessibility and BRD9 binding upon SMARCB1 rescue. Top-ranked classes are associated with translational control, suggesting that loss of SMARCB1 facilitates increased translation. (* indicates adj. p-value <0.05) B. GSEA of 307 downregulated genes that are associated with BRD9 degradation– dependent decreases in accessibility. These genes are associated with pro-migration gene expression programs. (* indicates adj. p-value <0.05). C. Representative phase-contrast microscopy images from 3 biological replicates of a 24 h scratch migration assay performed after 6 days of 100-nm dBRD9-A or DMSO treatment of GFP-induced G401 cells. White lines indicate the scratches generated at 0 h and the edges of cell migration zones at 24 h. The red line represents ∼350 µm. D. Quantification of relative migration was corrected for differences in proliferation measured by cell count. Relative migration values of dBRD9-A–treated cells were transformed by using a normalization factor of 1.25 to account for the decreased proliferative capacity of dBRD9-A-treated cells. 2-tailed, paired-measures *t*-test p=0.0234.

Given that SMARCB1-deficient cancers require BRD9, we sought to determine how the 307 BRD9-dependent genes might be supporting the cancer phenotype resulting in BRD9 essentiality. We used GSEA to evaluate gene expression programs associated with BRD9-dependent maintenance of accessibility in G401 cells (Figure 1 D, G). The 307 genes that were overexpressed due to BRD9-dependent genomic accessibility were significantly enriched in categories impacting cell migration (Figure 4B). Specifically, we identified BRD9-dependent genes involved in extracellular matrix organization and positive regulation of motility and chemotaxis (Figure 4B). The remaining 866 genes downregulated after BRD9 degradation were not associated with BRD9-dependent changes in accessibility, so their downregulation likely represented secondary effects. These genes were enriched in categories involved in cell cycle and metabolism (Supplemental Figure 3 E-F), perhaps reflecting the cell-cycle arrest and reduced proliferation that occurs following prolonged loss of BRD9 (Brien et al., 2018; Michel et al., 2018; X. Wang et al., 2019).

Because our data suggested that BRD9 loss may compromise migratory potential, we asked whether BRD9 degradation affected the migration of G401 cells. We performed a scratch migration assay and measured the relative migration of DMSO-vs. dBRD9-A treated G401 cells. To account for any differences in proliferation due to BRD9-degradation, we normalized to the difference in cell number after the 24-h scratch assay. We found that BRD9 degradation impaired migration by 22% (Figures 4C, D), providing a potential link to the essential role of BRD9 in SMARCB1-deficient cells.

## Discussion

There are several families of SWI/SNF complexes – each containing several mutually shared subunits but also family-specific subunits. The complexes all bind to active regions of chromatin, but the interrelationship of the families is poorly understood. Malignant rhabdoid tumors present a unique opportunity to evaluate the relationship between families because this cancer type is driven by loss of SMARCB1, which is present in two of the three families but not in the third, BRD9-containing ncBAF. The biological importance of the relationship is highlighted by the finding from multiple groups that BRD9 constitutes a specific dependency in SMARCB1-deficient cancer cell lines. Indeed, dependency upon BRD9 in the Cancer Dependency Map is markedly enriched for SMARCB1-deficient cell lines. This includes malignant rhabdoid tumors, ATRT (the nomenclature given to RT that arise in the brain), and synovial sarcomas in which fusion of the SS18 subunit of SWI/SNF to SSX displaces SMARCB1 from SWI/SNF complexes (Brien et al., 2018; J. Li et al., 2021; McBride et al., 2018). Because of such data, BRD9 inhibitors are now in clinical development. We, therefore, used the RT model to investigate the role of BRD9 in the presence and absence of SMARCB1.

We find that SMARCB1-containing SWI/SNF complexes can bind independently from BRD9-containing ncBAF as well as facilitate binding of BRD9-containing ncBAF at many sites. SMARCB1-BAF and BRD9-BAF cooperatively bind enhancers; however, at most sites, only SMARCB1-containing complexes serve a measurable role in modulation of the chromatin landscape and transcription. Such relationships might suggest that BRD9-containing complexes would largely be inactivated by SMARCB1 loss. However, that is not the case as BRD9 becomes a specific dependency in the absence of SMARCB1. This finding drew our attention to the smaller but significant number of sites at which the loss of SMARCB1 results in gain of genomic accessibility and gene expression. Some of these sites show strong BRD9 binding and BRD9-dependent hyper-accessibility and are associated with genes that are overexpressed in the absence of SMARCB1. These sites likely underlie the unique role of BRD9 as a dependency in the absence of SMARCB1.

While it is not entirely clear whether specific pathways are activated by BRD9 in the absence of SMARCB1, we did identify upregulation of genes supporting cell migration and found that loss of BRD9, in addition to triggering cell death, may impair migration of RT cells. However, it may be challenging to distinguish reduced migration derived from specific impairment of gene expression pathways from reduced migration due to toxic effects of BRD9 loss as senescent or dying cells may also display reduced migration. Collectively, our data suggest that at certain enhancers, the binding of SMARCB1-BAF precludes binding of BRD9/ncBAF such that loss of SMARCB1 enables ncBAF binding and upregulation of genes that are essential for the survival of SMARCB1-deficient cancers.

## Materials and Methods

### Cell culture and drug treatment

G401 cells were purchased from ATCC (Manassas, VA) and grown in McCoy’s 5A base medium supplemented with 1% GlutaMax (Gibco, Waltham, MA) and TET-approved 10% FBS (BioWest, Bradenton, FL). Cells were regularly verified to be mycoplasma-negative by using PCR-based MycoScope. Cell identity was confirmed via STR profiling. Cells were treated with 100 nm dBRD9-A (Tocris, Bristol, UK #6943) for 24 h to degrade BRD9.

### Lentivirus

CMV-TET3G-BSD, TRE-V5-SMARCB1-PURO, and TRE-GFP-PURO lentiviral vectors were purchased from VectorBuilder (Chicago, IL). Lentivirus was produced by the St. Jude Vector Core in 293T cells by using third-generation transfection reagents. Lentivirus was purified by performing ion exchange, diafiltration, and 0.22-µm sterile filtration.

### Generation of SMARCB1-inducible G401 cells

G401 cells expressing TET3G regulatory protein with doxycycline-inducible SMARCB1 or GFP were generated as described in (Drosos et al., 2022).

### Western blotting

Whole-cell lysates of G401 cells simultaneously induced to express SMARCB1 or GFP and treated with 100 nm dBRD9-A or DMSO for 24 h were harvested in RIPA (25 mM Tris-HCl. pH7.6, 150 mM NaCl, 1% NP-40, 0.1% SDS, 0.5% Sodium Deoxycholate) containing 1X protease inhibitors for 10’ on ice, vortexing regularly. Lysates were boiled in sample buffer containing lithium dodecyl sulfate and 10% β-mercapto ethanol (BME). Protein samples were resolved on Invitrogen (Waltham, MA) 4-12% precast gels in MOPS buffer. Proteins were transferred to nitrocellulose membrane overnight, and blots were probed for SMARCC1 (Cell Signaling, Danvers, MA #11956), BRD9 (Cell Signaling #58906), and SMARCB1 (Cell Signaling #91735) diluted in 3% non-fat milk in TBS with 0.05% Tween-20 overnight. Relevant secondary antibodies were applied for one hour at room temperature. Images were captured by using the BioRad ChemiDoc touch imager.

### Glycerol gradient density sedimentation assay

Cells were harvested in homogenization buffer (HB) (10mM HEPES, 10mM KCl, 1.5 mM MgCl, 0.1mM EGTA, 1:100 protease inhibitor cocktail and passed through a 26G needle 7 times to separate the nuclei from the cytoplasm. Nuclei were washed once in fresh HB and then lysed in buffer K (10 mM HEPES, 3 mM MgCl, 100 mM KCl, 300 mM ammonium sulphate, 0.5 mM EDTA, 10% Glycerol) on ice for 20 minutes, vortexing every 2 minutes. Insoluble material was pelleted by centrifugation, and the nuclear lysate was transferred to a fresh tube. Ammonium sulphate was added at a ratio of 0.5 mg per µL. Nuclear lysate was then precipitated for 20 minutes rotating at 4^°^C. Precipitated proteins were separated from supernatant by centrifugation, and the pellet containing nuclear protein was resuspended in buffer HK (100 mM Tris HCl pH 7.4, 150 mM NaCl, 1 mM EDTA, 1 mM EGTA, 1% Triton-X100). Then, 1 mg of nuclear extract was deposited over a 14-mL 10-30% glycerol gradient prepared with HK buffer in a 14× 95-mm centrifuge tube (Beckman Coulter, Brea, CA Cat. # 344060) by using the Biocomp Gradient Station (Fredericton, NB, Canada, Model 153). Gradients were placed in a SW-40 Ti 40000 RPM rotor (Beckman Coulter) and centrifuged (40000x RPM for 16 hours, 4°C). 20 fractions were collected per sample by using the Biocomp Gradient Station (Model 153) piston fractionator. Fractions were subjected to gel electrophoresis on BioRad (Hercules, CA) Criterion TGX stain-free gels and then western blotting analysis.

### ATAC-seq

ATAC-seq was performed by using the Active Motif ATAC-seq kit (Carlsbad, CA 53150) exactly as prescribed by the manufacturer. G401 cells were simultaneously induced to express GFP or SMARCB1 and treated with 100 nm dBRD9-A or DMSO for 24 h. Then, cells (100K) were washed with PBS, and nuclei were extracted by using the provided ATAC-lysis buffer. Nuclei were suspended in tagmentation master mix (proprietary), and tagmentation was carried out at 37°C for 30 minutes on a thermomixer agitating at 800 rpm. Next, 5x DNA purification binding buffer and 5 µL 3M sodium acetate were added to each sample before it was passed through a DNA purification column. Columns were washed once and dried before DNA was eluted in 35 µL of DNA purification elution buffer. Libraries were prepared from the total eluate of purified DNA by using the included library preparation reagents; >100M 100-bp paired-end reads were generated per sample by using NovaSeq 6000.

### RNA-seq

RNA was harvested by using the Qiagen RNeasy plus mini kit (Germantown, MD, 74034) according to the manufacturer’s instructions. Briefly, cells were lifted from a 6-well dish in RLT+ 1% BME buffer. ERCC spike-in controls (Ambion, Austin, TX) were added on a per-cell basis for downstream normalization. Cells in RLT were homogenized by passing them through a 20-g needle 5 times. Cell homogenate was passed through DNA elimination columns before RNA was purified using the columns and buffers provided. RNA was quantified by using the Quant-iT RiboGreen assay (Life Technologies, Carlsbad, CA) and quality-checked via a 2100 Bioanalyzer RNA 6000 Nano Assay (Agilent, Santa Clara, CA) or the LabChip RNA Pico Sensitivity Assay (PerkinElmer, Waltham, MA) before library generation. The libraries were prepared from total RNA by using the TruSeq Stranded mRNA Library Prep kit according to the manufacturer’s instructions (Illumina, San Diego, CA). The libraries were analyzed for insert size distribution by using a 2100 BioAnalyzer High Sensitivity kit (Agilent). Then, ∼100M, 100-bp, paired-end reads were generated by using NovaSeq 6000 for 3 biological replicates of each sample.

### Mnase ChIP-seq

Mnase ChIP was performed by using the SimpleChIP Enzymatic Chromatin IP Kit (Cell Signaling 9003). Cells were fixed for 10’, rocking at room temperature in 1% methanol-free paraformaldehyde diluted in PBS. Fixation was stopped by using 1x glycine solution. Digestion was carried out with 0.25 µL Mnase per million cells for 20’ at 37°C. Histone modification ChIPs were performed by using 10 µg of chromatin each incubated with the recommended amount of H3K27Ac (Cell Signaling #8173, 5ul), H3K4me1 (Cell Signaling #5326, 10 µL), and H3K4me3 (Cell Signaling #9751, 10 µL), respectively. SMARCA4 ChIP was performed using 20 µg of chromatin incubated with 3 µg of SMARCA4 antibody (ABCAM, Cambridge, UK 110641). Digested chromatin and antibody were incubated overnight. The following day, 30 µL protein-G beads were added for 2 h and then washed for 3 minutes with low-salt buffer (20m M Tris-HCl pH 8.0, 150 mM NaCl, 2 mM EDTA pH 8.0, 1% Triton X-100, 0.1% SDS), high-salt buffer (20mM Tris-HCl pH 8.0, 500 mM NaCl, 2 mM EDTA pH 8.0, 1% Triton X-100, 0.1% SDS), and then LiCl buffer (10 mM Tris-HCl pH 8.0, 250 mM LiCl, 1 mM EDTA pH 8.0, 1% NP-40, 1% sodium deoxycholate), one time each before a final TE pH 8.0 buffer (Invitrogen AM9849) rinse. Elution was carried out at 65°C for 30 minutes by using the included elution buffer.

Decrosslinking and protein digestion were performed at 65°C for 2 h in 200 µM NaCl and 40 µg Proteinase K. Decrosslinked samples were mixed with 5x DNA-binding buffer and passed through the DNA purification columns. The columns were washed with DNA wash buffer and dried before DNA elution in 51 µL of DNA elution buffer. DNA was quantified using PicoGreen fluorescent assay (Molecular Probes, Eugene, OR, P-7581). Libraries were prepared by using the KAPA HyperPrep kit (Roche, Basel, Switzerland, # 7962363001) according to the manufacturer’s specifications, and library quality was determined by using the TapeStation (Agilent) with D1000 screentape. Then, >100k 50-bp paired-end reads per sample were generated on the NovaSeq 6000.

### CUT & RUN

CUT & RUN was performed using the Epicypher (Durham, NC) Cutana Cut&Run Kit v2.0 according to the manufacturer’s protocol as reported (Drosos et al. 2022). Briefly, permeabilization was achieved using 0.08% digitonin (Cell Signaling Technology #16359). For each sample 0.5 million cells were used. SMARCB1 (Cell Signaling Technologies #91735, 1ul) and BRD9 (ABCAM ab137245 0.5ul) antibodies were incubated with bead-bound, permeabilized cells overnight. CUT & RUN DNA was quantified by performing a Qubit high-sensitivity fluorescent assay (Q32851). Next, 2-5 ng of DNA was used for library preparation using the paired-end KAPA Hyper library kit. Library concentration and size distribution were assessed by using the Agilent TapeStation and D1000 high-sensitivity Screentape. Then, >100M 75-bp paired-end reads for two biological replicates of SMARCB1 and >10M 75-bp, paired-end reads for three biological replicates of BRD9 were generated by using the NovaSeq 6000 sequencer.

### ATAC-seq analysis

Quality trimmed reads are mapped to the HG19 reference genome by BWA (v0.7.12-r1039, (H. Li & Durbin, 2009)) and converted to bam file by SAMtools (v1.2, (H. Li et al., 2009)). biobambam2 (v2.0.87, (Tischler & Leonard, 2014)) was used to mark duplicated reads. Nucleosome-free fragments and nucleosomal fragments were extracted by bedtools (v2.24.0, (Quinlan & Hall, 2010)). Narrow peaks were called by MACS2 (v2.1.1.20160309, (Zhang et al., 2008)) on nucleosome-free reads (fragment size < 100 bp) by using the false-positive rate corrected p-value 0.05. Differential accessibility was called on nucleosome-free regions identified from 3 biological replicates of each sample by using R packages diffbind (v3.6.2) and deseq2 (v3.15). Background normalization was utilized, and nucleosome free regions were only considered if identified in two of the three replicates. Averaged bedgraphs were generated for ATAC-seq nucleosomal and nucleosome-free regions by (1) combining replicates UCSC-bigWigMerge, (2) sorting combined bedgraphs (LC_COLLATE=C sort -k1,1 -k2,2n), and (3) averaging the 4^th^ bedgraph column based on the number of input replicates using awk.

### RNA-seq analysis

The RNA-seq reads were mapped to the human GRCh37-lite (HG19) reference genome via STAR (Dobin et al., 2013). Gene-level counts were quantified by RSEM (v1.3.1) with GENCODE version 19 Ensembl 74 annotation. The read counts were further normalized by using the R package DESeq2. Differential gene expression was called by using the R package DESeq2. Gene ontology enrichment analyses of up- or downregulated gene lists were performed by using the GO Enrichment Analysis by Enrichr (https://maayanlab.cloud/Enrichr/).

### CUT & RUN analysis

Paired-end reads of 75 bp were mapped to the human genome hg19 (GRCh37-lite) and the *Escherichia coli* genome r46 (Ensembl r46 ASM584v2) hybrid by BWA (version 0.7.12-r1039, default parameter (H. Li & Durbin, 2009)). The duplicated reads were then marked by biobambam2 (version 2.0.87 (Tischler & Leonard, 2014)). After the duplicated reads were marked, bam files were split by either the human bam file or the *Escherichia coli* spike-in bam file. For the spike-in bam files, we counted the nonduplicated reads with SAMtools. For the human bam files only, the uniquely, properly paired reads were extracted by SAMtools, (version 1.2, parameter -q 1 -F 1804 (H. Li et al., 2009)) sorted based on read name, and converted to BEDPE format by using bedtools. Fragment sizes <2000 bp were extracted, and the center 80 bp of each fragment were used to generate normalized (to 10 million fragments) bigwig tracks for visualization. Peaks were called against IgG control for BRD9 and GFP-induced cells assayed with the same antibody for SMARCB1. Peaks were identified by using Easeq (Adaptive Local Thresholding, v1.111, (Lerdrup et al., 2016)) with a p-value and FDR cutoff of 1×10^-4^ and a log2(fold-change) over control cutoff of 0.5. SMARCB1 peaks used in this study are peaks called in both biological replicates. For BRD9 peaks, differential binding analysis was accomplished using only peaks occurring in at least 2 of the 3 biological replicates for a given sample by using R packages diffbind and edger. Differential binding was determined using FDR and Log2(fold-change) cutoffs of 0.05 and 0.5, respectively. Averaged bedgraphs were generated for CUT & RUN by (1) combining replicates UCSC-bigWigMerge, (2) sorting combined bedgraphs (LC_COLLATE=C sort -k1,1 -k2,2n), and (3) averaging the 4^th^ bedgraph column based on number of input replicates using awk.

### Mnase-ChIP seq analysis

Single-end reads of 50 bp were mapped to the human genome hg19 (GRCh37-lite) by BWA (version 0.7.12-r1039, default parameter (H. Li & Durbin, 2009)). The duplicated reads were then marked by biobambam2 (version 2.0.87 (Tischler & Leonard, 2014)). H3K27Ac peaks were called against the input sample by using Easeq (Adaptive Local Thresholding) with a p-value and FDR cutoff of 1×10^-4^ and a log2(fold-change) over control cutoff of 0.5. Averaged bigwigs were generated for ChIP or RNA-Seq by (1) combining replicates UCSC-bigWigMerge, (2) sorting combined bedgraphs (LC_COLLATE=C sort -k1,1 -k2,2n), and (3) averaging the 4^th^ bedgraph column on the basis of the number of input replicates using awk.

### Identification of traditional and super enhancers

Two biological replicates of H3K27Ac data per sample were used with Rank Ordering of Super Enhancers (ROSE) to identify traditional and super enhancers in GFP- and SMARCB1-induced G401 cells. Regions within 2.5 kb of a TSS were excluded. Enhancers reproducibly called in both samples were used for annotations.

### Binding and Expression Target Analysis (BETA)

BETA was performed to associate SMARCB1 and BRD9 bound increases or decreases in accessibility (FDR < 0.05) identified by ATAC-seq with differentially expressed genes (adj. p < 0.05) identified between SMARCB1- and GFP-induced G401 cells. BETA was also used to associate BRD9-degradation–dependent increases or decreases in accessibility (FDR < 0.05) identified by ATAC-seq in GFP-induced G401 cells with differentially expressed genes (adj. p < 0.05) between dBRD9-A and DMSO treated, GFP-induced G401 cells.

### Scratch migration assay

GFP-inducible G401 cells were treated with DMSO or 100 nM dBRD9-A for 5 days before 1 million cells of each were plated in a 6-well dish in their respective conditions (DMSO vs. dBRD9-A). On the 6^th^ Day, three ∼700-µM scratches were made per well by using a sterile P1000 pipette tip. Baseline, 0-h scratches with reference points were imaged by using phase microscopy with an Evos imaging system. Cells were maintained in their respective conditions overnight and imaged again at 24 h. Cell migration distance was measured by counting pixels between the leading edges of the scratches, averaging 5 locations on each scratch. Three scratches per well were measured (technical replicates) from 3 independently grown and treated cell cultures (biological replicates) each. After 24-h imaging, cells were harvested and counted to account for differences in proliferation over the 24-h assay. The percentage of dBRD9-A–treated cell migration was multiplied by a normalization factor of 1.25 to account for the measured difference in cell number between the two conditions. The two-tailed, students *t*-test was used to assess the difference between the normalized values from the two conditions.

## Data Availability

All next-generation sequencing datasets generated in association with this work have been deposited in the Gene Expression Omnibus (GEO) under accession number GSE 210636.

## Acknowledgements

Thanks to all the Roberts’ lab members for intellectually engaging conversations. This work was supported by the National Cancer Institute (NCI) R01 CA113794 and R01 CA172152 to C.W.M.R, CURE AT/RT Now to C.W.M.R, Garrett B. Smith Foundation to C.W.M.R, St. Jude Children’s Research Hospital Collaborative Research Consortium on Chromatin Regulation in Pediatric Cancer to C.W.M.R. HAM is funded by the St. Jude Graduate School of Biomedical Sciences. We thank Hartwell Center Core for library preparation and sequencing. We thank St Jude Center for Applied Bioinformatics (CAB) for help with alignment of ATAC-, RNA-, and ChIP-seq, and CUT & RUN reads. We thank the Scientific Editing department for their helpful suggestions. St. Jude cores are supported via Cancer Center support grant (NCI CCSG 2 P30 CA021765) and ALSAC of St. Jude Children’s Research Hospital.

## Author Details

### Robert J Mobley

- Division of Molecular Oncology, Department of Oncology, St. Jude Children’s Research Hospital, Memphis, TN, USA **Contribution**: Conceptualization, Methodology, Validation, Formal analysis, Investigation, Writing-original draft, Writing-review and editing, Visualization, Supervision, Project Administration **Competing interests:** No competing interests declared

### Jacquelyn A Myers

- Division of Molecular Oncology, Department of Oncology, St. Jude Children’s Research Hospital, Memphis, TN, USA **Contribution:** Software, Formal analysis, Data curation **Competing interests:** No competing interests declared

### Kendall M Wills

- Division of Molecular Oncology, Department of Oncology, St. Jude Children’s Research Hospital, Memphis, TN, USA **Contribution:** Validation, Investigation, Writing-review and editing **Competing interests:** No competing interests declared

### Hayden A Malone

- Division of Molecular Oncology, Department of Oncology, St. Jude Children’s Research Hospital, Memphis, TN, USA
- St. Jude Graduate School of Biomedical Sciences, St. Jude Children’s Research Hospital, Memphis, TN, USA **Contribution:** Validation, Investigation, Writing-review and editing **Competing interests:** No competing interests declared

### Trishabelle I Manzano

- Division of Molecular Oncology, Department of Oncology, St. Jude Children’s Research Hospital, Memphis, TN, USA **Contribution:** Validation, Investigation **Competing interests:** No competing interests declared

### Janet F Partridge

- Division of Molecular Oncology, Department of Oncology, St. Jude Children’s Research Hospital, Memphis, TN, USA **Contribution:** Conceptualization, Writing-original draft, Writing-review and editing, Supervision, Project Administration **Competing interests:** No competing interests declared

### Charles WM Roberts

- Division of Molecular Oncology, Department of Oncology, St. Jude Children’s Research Hospital, Memphis, TN, USA **Contribution:** Conceptualization, Writing-review and editing, Supervision, Project Administration, Funding acquisition **Competing interests:** No competing interests declared

## Notes

https://www.ncbi.nlm.nih.gov/geo/query/acc.cgi?acc=GSE210636

### Competing Interest Statement

The authors have declared no competing interest.

