## Supplemental figures for "Functional interplay between SWI/SNF complexes underlies BRD9 dependency in SMARCB1-mutant cancers"

Supplemental Figures 1-3

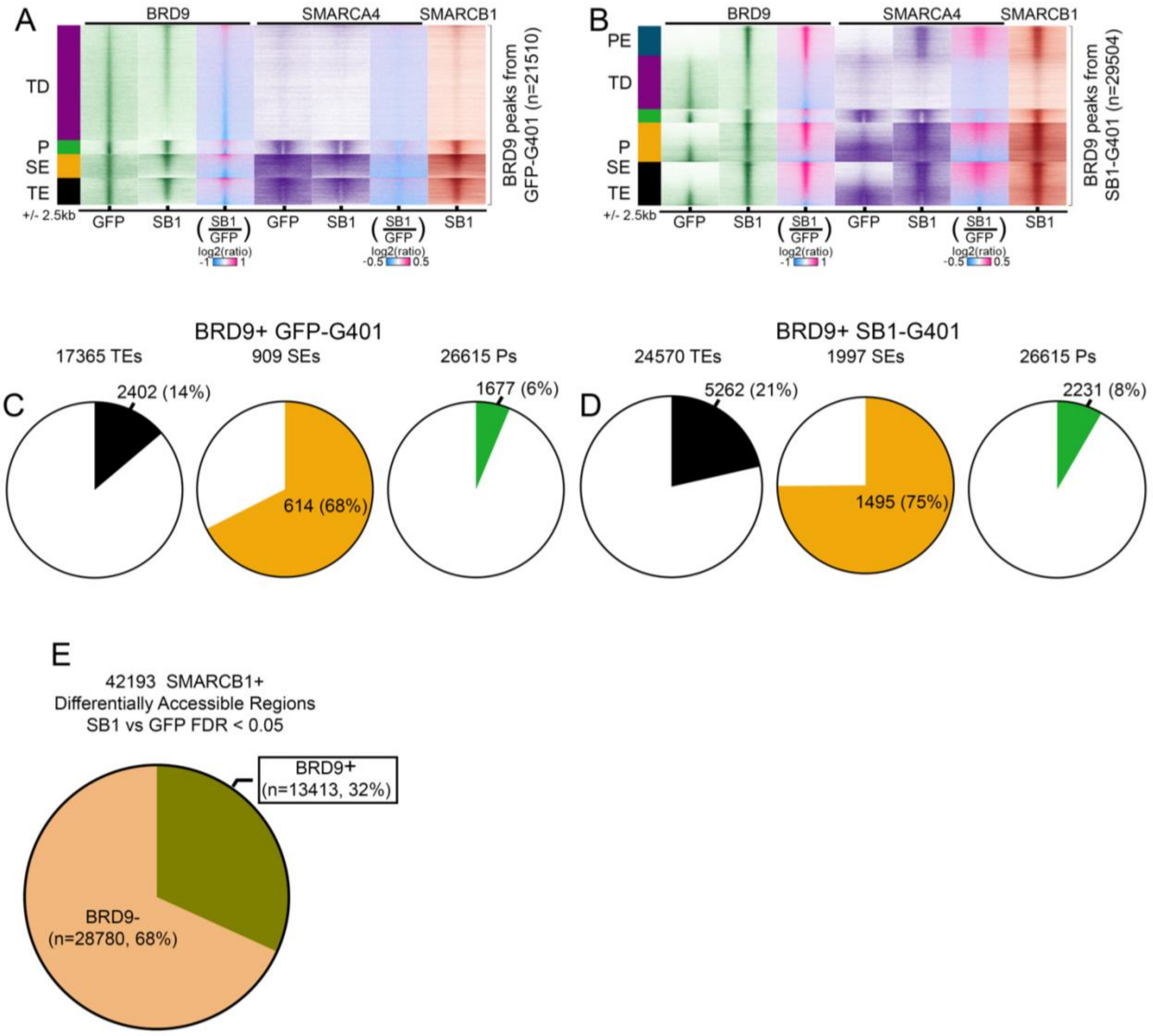

**Supplemental figure 1 supporting Figure 2:** Association of BRD9 genomic binding with gene regulatory elements.

A-B. Genomic heatmaps showing BRD9, SMARCA4, and SMARCB1 binding for the indicated samples at BRD9 peaks called in control (GFP-induced, A) and SMARCB1-induced (C) G401 RT cells. Data are sorted by association with traditional enhancers (TEs) and super enhancers (SEs), promoters (Ps), primed enhancers (PEs), and/or TSS-distal sites (TDs). Within these clusters, sites are sorted by fold-change of BRD9 enrichment. BRD9 CUT & RUN data are the average of 3 biological replicates. SMARCB1 CUT & Run and SMARCA4 ChIP data are the average of 2 biological replicates.

C-D. Circle graphs show the proportion of all TEs, SEs, and Ps at which BRD9 is enriched in GFP-induced (A) or SMARCB1-induced (C) G401 cells.

E. Circle graph of SMARCB1-bound regions (CUT & RUN, n=2) that have differential ATAC-seq accessibility between SMARCB1-induced and GFP-induced G401 cells (FDR<0.05, n=3). These sites either bind (olive) or do not bind (orange) BRD9 in CUT & RUN (n=3) from GFP or SMARCB1-induced G401 cells.

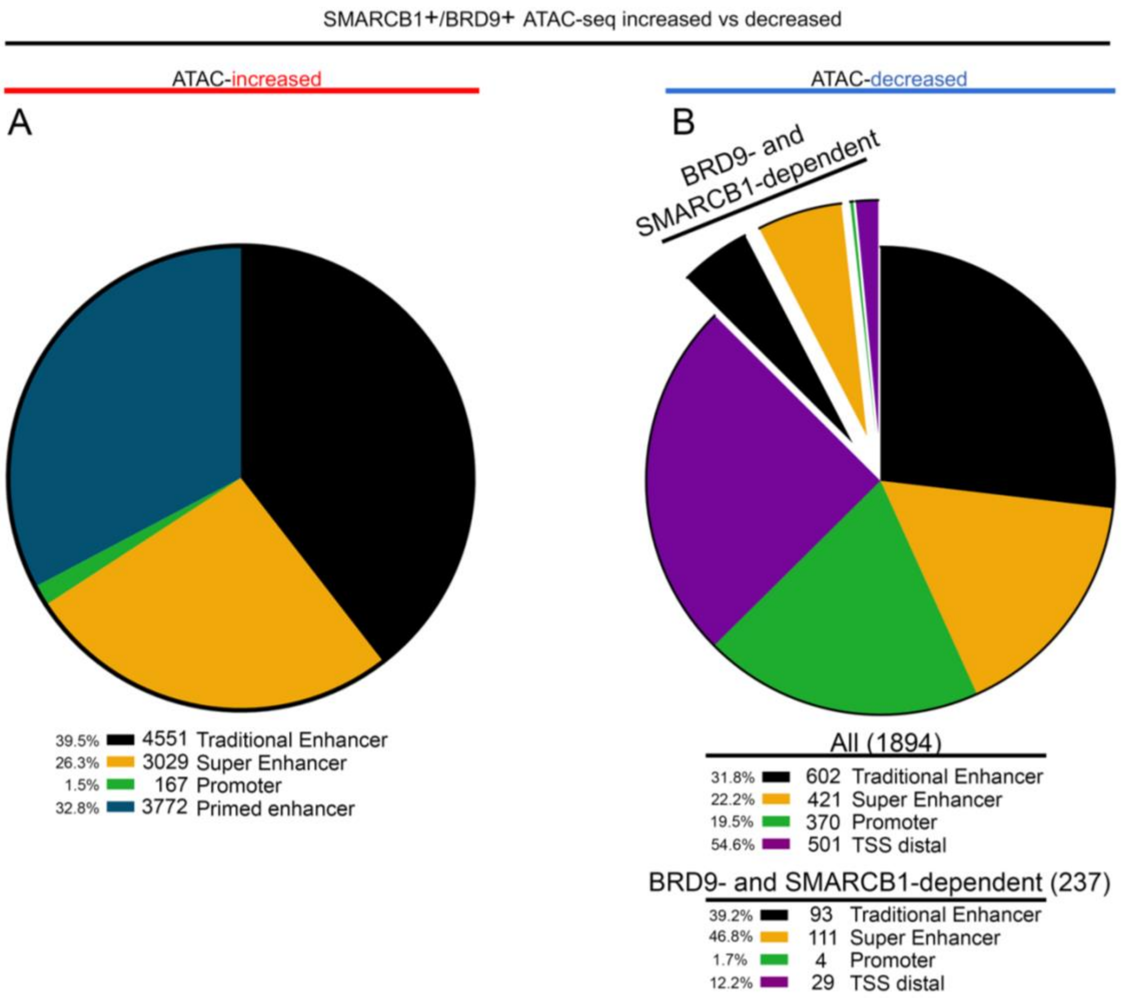


**Supplemental figure 2 supporting Figure 3:** SMARCB1- and BRD9-bound differentially accessible regions show distinct association with gene regulatory elements.

A-B. Circle graphs show distribution of SMARCB1- and BRD9-bound differentially accessible regions that are increased (A) or decreased (B) upon SMARCB1-rescue within the indicated gene regulatory elements. The 237 regions that show loss of accessibility upon SMARCB1-rescue or BRD9 degradation in G401 cells are expanded and annotated as “BRD9- and SMARCB1-dependent”.

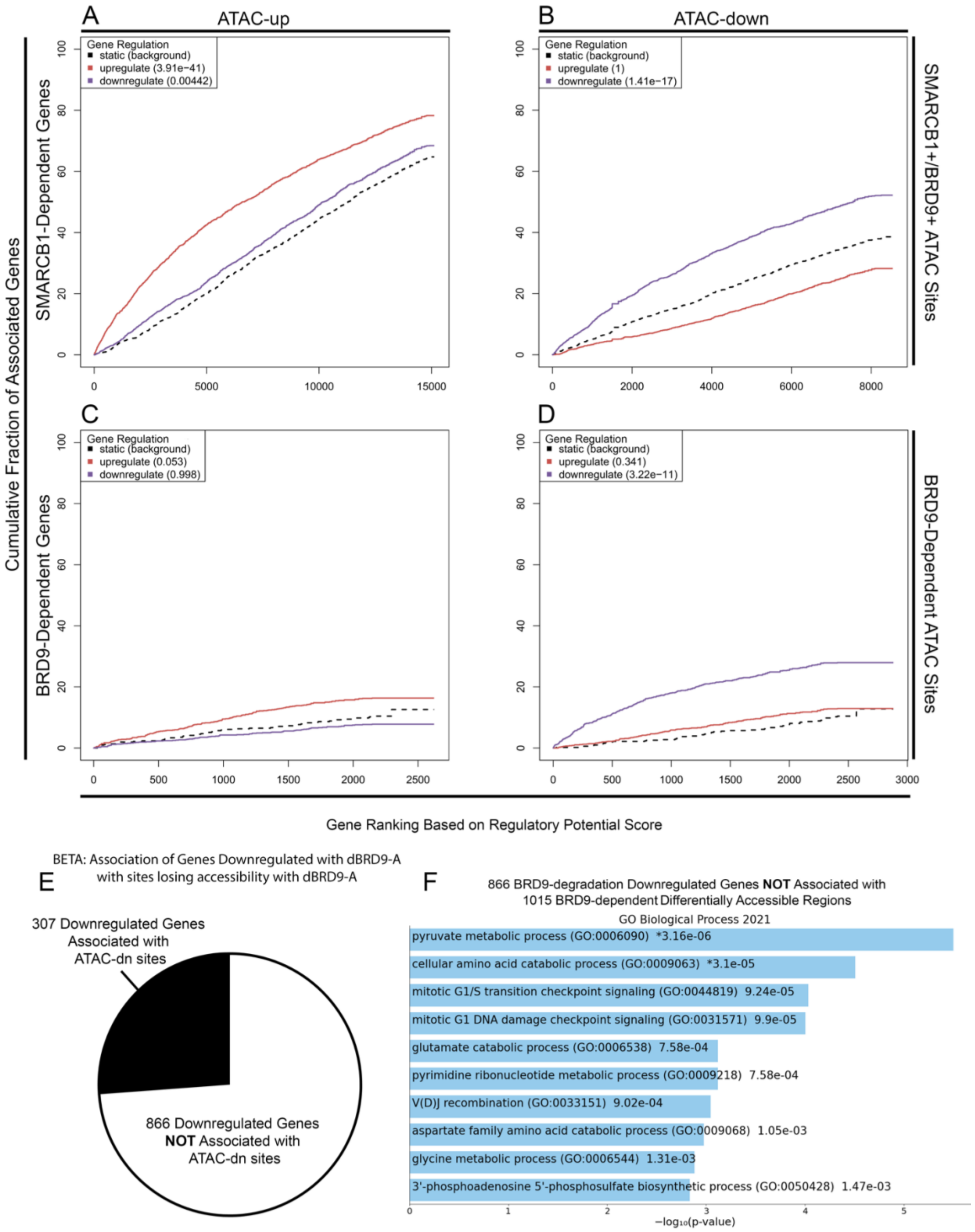
A
ATAC-up
B
ATAC-down
Gene Regulation
static (background)
upregulate (3.91e−41)
downregulate (0.00442)
Gene Regulation
static (background)
upregulate (1)
downregulate (1.41e−17)
C
D
Gene Regulation
static (background)
upregulate (0.053)
downregulate (0.998)
Gene Regulation
static (background)
upregulate (0.341)
downregulate (3.22e−11)
Cumulative Fraction of Associated Genes
SMARCB1-Dependent Genes
BRD9-Dependent Genes
SMARCB1+/BRD9+ ATAC Sites
BRD9-Dependent ATAC Sites
Gene Ranking Based on Regulatory Potential Score
BETA: Association of Genes Downregulated with dBRD9-A
with sites losing accessibility with dBRD9-A
E
307 Downregulated Genes
Associated with
ATAC-dn sites
866 Downregulated Genes
NOT Associated with
ATAC-dn sites
F
866 BRD9-degradation Downregulated Genes NOT Associated with
1015 BRD9-dependent Differentially Accessible Regions
GO Biological Process 2021
pyruvate metabolic process (GO:0006090) *3.16e-06
cellular amino acid catabolic process (GO:0009063) *3.1e-05
mitotic G1/S transition checkpoint signaling (GO:0044819) 9.24e-05
mitotic G1 DNA damage checkpoint signaling (GO:0031571) 9.9e-05
glutamate catabolic process (GO:0006538) 7.58e-04
pyrimidine ribonucleotide metabolic process (GO:0009218) 7.58e-04
V(D)J recombination (GO:0033151) 9.02e-04
aspartate family amino acid catabolic process (GO:0009068) 1.05e-03
glycine metabolic process (GO:0006544) 1.31e-03
3'-phosphoadenosine 5'-phosphosulfate biosynthetic process (GO:0050428) 1.47e-03
−log₁₀(p-value)

**Supplemental figure 3 supporting Figure 4:** Binding and Expression Target Analysis (BETA) identifies gene regulatory roles associated with accessible regions losing SMARCB1 and regions with BRD9-dependent accessibility in RT.

A. BETA cumulative distribution plot of $SMARCB1^{+}/BRD9^{+}$ ATAC-up sites associated with SMARCB1-dependent changes in gene expression within a 200-kb window.

B. BETA cumulative distribution plot of $SMARCB1^{+}/BRD9^{+}$ ATAC-down sites associated with SMARCB1-dependent changes in gene expression within a 200-kb window.

C. BETA cumulative distribution plot of BRD9-dependent ATAC-up sites associated with BRD9 degradation–dependent changes in gene expression within a 200-kb window.

D. BETA cumulative distribution plot of BRD9-dependent ATAC-down sites associated with BRD9 degradation–dependent changes in gene expression within a 200-kb window.

E. Circle graph showing all genes downregulated upon BRD9-degradation depending on their association or lack of association with BRD9-dependent accessibility.

F. GSEA of genes downregulated upon BRD9 degradation that are not associated with BRD9-dependent changes in accessibility. These effects are likely to be secondary/indirect. (* indicates adj. p-value <0.05)
